# When Age Tips the Balance: a Dual Mechanism Affecting Hemispheric Specialization for Language

**DOI:** 10.1101/2023.12.04.569978

**Authors:** Elise Roger, Loïc Labache, Noah Hamlin, Jordanna Kruse, Monica Baciu, Gaelle E. Doucet

**Affiliations:** Institut Universitaire de Gériatrie de Montréal, Communication and Aging Lab, Montreal, Quebec, Canada; Faculty of Medicine, University of Montreal, Montreal, Quebec, Canada; Univ. Grenoble Alpes, Univ. Savoie Mont Blanc, CNRS, LPNC, 38000 Grenoble, France; Department of Psychology, Yale University, New Haven, CT, 06520, US; Department of Psychiatry, Brain Health Institute, Rutgers University, Piscataway, NJ, 08854, US; Institute for Human Neuroscience, Boys Town National Research Hospital, Omaha, NE, 68010, US; Department of Pharmacology and Neuroscience, Creighton University School of Medicine, Omaha, NE, 68178, US; Center for Pediatric Brain Health, Boys Town National Research Hospital, Omaha, NE, 68178, US

## Abstract

Aging leads to neuroadaptations, often reducing specificity in functional brain responses. However, the extent to which functional specialization of brain hemispheres in the Language-and-Memory Network regions changes with adulthood remains unclear. In a cohort of healthy adults, we provide evidence that aging is linked to shifts in the cortical organization’s lateralization, unveiling two age-related asymmetry patterns. The first pattern indicates a lateralization shift in language regions, with aging rendering language functions more bilateral. The second pattern reveals increased specialization in memory regions within the left hemisphere as we age. Manifesting around midlife, these shifts correlate with declines in language production performance. Our findings offer new insights into how functional brain asymmetries impact language proficiency and underscore brain plasticity in aging, providing a dynamic view of the aging brain’s functional architecture.

---

**P**aul Pierre Broca challenged the prevailing 19th-century belief that the brain was organized holistically and symmetrically, providing evidence to support the idea that a prominently lateralized brain is a crucial characteristic for effectively carrying out certain cognitive functions, particularly language processing (“*We speak with the left hemisphere*,” Broca, 1865, p. 384).

Patterns of hemispheric specialization and interaction of brain networks are complex, developmental, learning-dependent, and dynamic^1^. From the earliest stages of development, human beings demonstrate behavioral and brain asymmetries – as early as ten weeks prenatal^2^ and 26 gestational weeks for perisylvian regions^3^ – which become increasingly perceptible functionally and anatomically during infancy. Asymmetries in neural networks take effect at different times during ontogeny, and almost all cortical brain regions show significant left-right asymmetries in adulthood^4^. Language-related regions show covariate developmental trajectories^5^ and develop more slowly in the left hemisphere (LH) than in the right^6^. A notable right hemisphere (RH) language activation pattern in young children typically diminishes with age and becomes strongly left lateralized for most adults^7^. Examining the language connectome in adult populations and its organization across several language tasks reveals a pronounced left-hemispheric dominance in the central perisylvian network, which specializes in processing auditory-verbal stimuli^8^. This dominance of functional connectivity in the left hemisphere (LH) for the “core” language network has been consistently observed^9–12^.

However, language processing requires the involvement of a wider brain network, encompassing the core perisylvian LH system but also several peripheral or marginal memory, executive, and sensorimotor systems^13^; also discussed as multiple language networks by Hagoort and colleagues^14,15^. The extended language connectome comprises many fine-tuned associative hubs^8^. It is sharpened to underpin effective communication by integrating the high-level, multimodal perceptual and cognitive information required for language processing^16^. This sophisticated processing system is thus extremely powerful, yet it is also susceptible to vulnerabilities. Associative hubs are indeed highly prone to damage^17^, and ensuring the optimal function of language hubs in later life comes at a considerable cost^8,18–20^. It is now well-documented that functional connectivity and network dynamics remodel with age^21–24^. Importantly, older adults exhibit a default-executive coupling when engaged in demanding tasks, characterized by increased prefrontal involvement and reduced suppression of the Default Mode Network. In contrast, younger individuals adjust their functional responses by deactivating Default Mode Network regions when performing the same tasks^25–28^. Overall, age-related changes are characterized by reduced specificity, selectivity, and lateralization of functional brain networks^29^. Nevertheless, the trajectory of hemispheric specialization for language during aging, the underlying mechanisms involved, and their impact on cognition are still largely unclear and require further investigation.

Functional asymmetries can be investigated using intrinsic functional connectivity, which offers the advantage of abstracting from task-related variability associated with the nature and difficulty of specific tasks. Resting-state networks exhibit spatial patterns that correspond with the networks observed during specific cognitive tasks^30–32^ specific regions have been identified as predisposed to language processing at rest^33^. Moreover, lateralization measures in key language hubs, derived from resting-state data, can predict functional lateralization during task performance^34,35^. Furthermore, recent studies on functional brain architecture have reported that resting-state networks exhibit a hierarchical organization characterized by smooth spatial transitions or gradients^36,37^. The principal gradient (G1), explaining the most variance in whole-brain functional connectivity, aligns with established cortical hierarchies that progressively process complex or heteromodal information from sensory inputs^38^ (see also Chang and colleagues for natural language processing^39^). Interestingly, the brain hemispheres do not show an identical pattern of organization on G1^40^, revealing a notable asymmetry for heteromodal networks linked to higher-order cognitive functions^41,42^. Furthermore, a recent study showed that individuals exhibiting atypical language lateralization display corresponding hemispheric differences in macroscale functional gradient organization, making G1 a marker of hemispheric specialization for language^43^. Therefore, examining functional asymmetries within intricate networks, such as those supporting language processing, and how they change with age can bring a new perspective considering the fundamental underlying functional architecture.

Our study aimed to track age-related changes in hemispheric asymmetry within an extended language network, leveraging the macroscale functional gradient G1 derived from resting-state data. We opted for the Language-and-Memory Network (LMN) due to its comprehensive ability to capture the nuanced dynamics of language in conjunction with other cognitive processes ^44^. The LMN integrates regions specialized in language processing with areas concurrently involved in language and advanced cognitive functions, such as memory and executive processes. Importantly, these heteromodal regions may undergo significant functional changes with aging. To model the functional trajectories over an age range from 18 to 88 years, we applied the Generalized Additive Mixed Models (GAMMs) technique, which has been previously used in structural MRI studies^45,46^. This allowed us to classify Language-and-Memory Network regions based on their asymmetry patterns at rest throughout healthy aging. Furthermore, we also explored how these asymmetry changes were related to cognitive performance measured during various language tasks. To this end, we used Canonical Correlation Analyses (CCA) to assess how age impacted asymmetries in the language network across multimodal data, including anatomy, function, and cognitive performances.

## Results

### Evolution of Hemispheric Gradient Asymmetries

We investigated the evolution of the functional connectivity architecture asymmetry within an extended Language-and-Memory Network^44^ (Fig. 1) across the adult lifespan using anatomical and resting-state fMRI data acquired at 3T (*n*=728, aged 18 to 88), combining three databases (Camcan, Omaha, and Grenoble sample). Demographics are available in the Methods section (Database Demographics).

**Figure 1.**
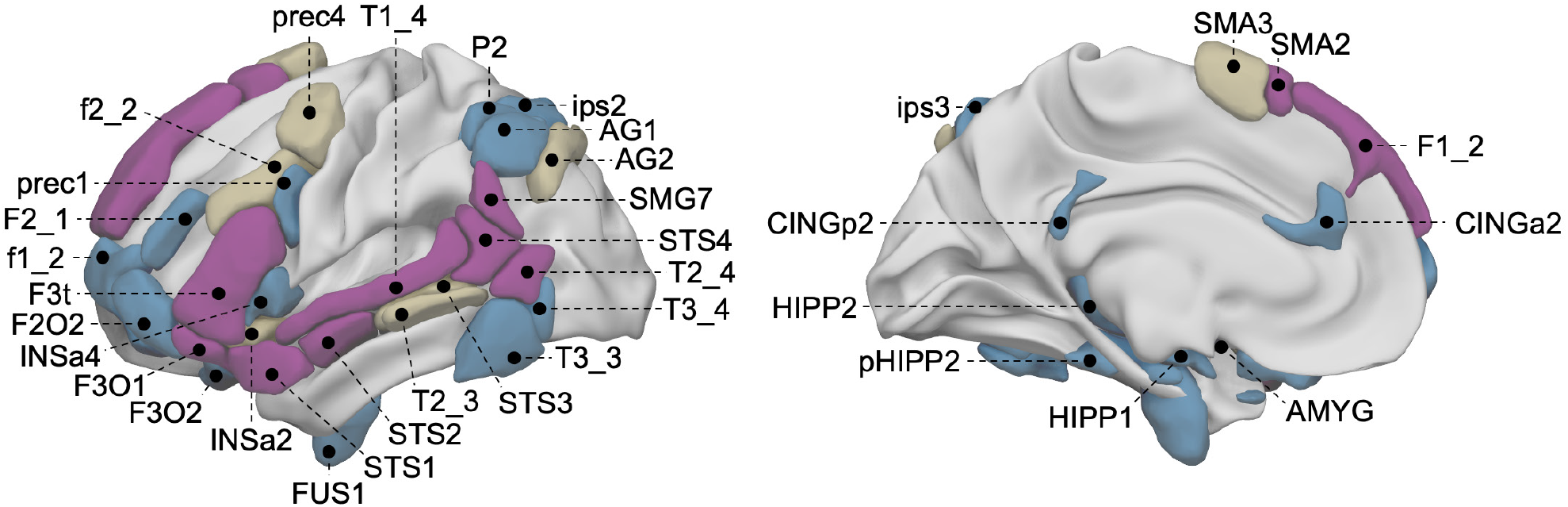
Locations of the 37 regions of the Language-and-Memory Network atlas^44^. On the left: lateral view of the left hemisphere. On the right: Medial view of the left hemisphere. The atlas is composed of 74 homotopic ROIs (37 in each hemisphere) reported by two task-fMRI studies, one cross-sectional study for language^9^, and one meta-analysis for memory^54^ and adapted to the Atlas of Intrinsic Connectivity of Homotopic Areas coordinates^48^. Regions are rendered onto the 3D anatomical templates of the white matter surface of the left hemisphere in the MNI space with Surf Ice software (https://www.nitrc.org/projects/surfice/). Color code: purple, regions involved in language; blue, regions involved in episodic memory (encoding and retrieval); brown, regions involved in both language and memory. The Anterior Insula (3) (INSa3) is not visible on this render. See Table 1 for the correspondences between the abbreviations and the full names of the Language-and-Memory Network regions.

As described by Labache and colleagues^47^, we took advantage of recent mathematical modeling of the cortex’s functional topography, as Margulies and colleagues proposed^37^. First, functional connectivity matrices (384×384 AICHA parcels^48^) across the full sample were decomposed into components that capture the maximum variance in connectivity. Consistent with prior work^37,49^, diffusion map embedding^50^ was used to reduce the dimensionality of the connectivity data through the nonlinear projection of the voxels into an embedding space. The resulting functional components or manifolds, termed gradients, are ordered by the variance they explain in the initial functional connectivity matrix. The present analysis focused on the first gradient accounting on average for 20% of the total variance in cortical connectivity (respectively 22% for the sample collected in Omaha, 20% for the CamCAN database, and 19% for the sample collected in Grenoble). In line with prior work^37,51–53^, one end of the principal gradient of connectivity was anchored in unimodal regions, while the other end encompassed broad expanses of the association cortex.

**Table 1.** List of the Language-and-Memory network atlas regions. Note: L=language; LM=language and memory; M=memory; MNI coordinates, in the left and right hemisphere, of regions (*X, Y, Z*) in *mm*; Total regions=74 (37 in each hemisphere).

| Abbreviation | Region | Function | MNI coordinates (left) |  |  | MNI coordinates (right) |  |  |
| --- | --- | --- | --- | --- | --- | --- | --- | --- |
|  |  |  | X (mm) | Y (mm) | Z (mm) | X (mm) | Y (mm) | Z (mm) |
| AG1 | Angular Gyrus (1) | M | -48 | -57 | 44 | 51 | -52 | 43 |
| AG2 | Angular Gyrus (2) | LM | -38 | -70 | 39 | 45 | -62 | 36 |
| AMYG | Amygdala (1) | M | -22 | 0 | -12 | 21 | 2 | -12 |
| CINGa2 | Anterior Cingulate Gyrus (2) | M | -7 | 34 | 22 | 7 | 33 | 23 |
| CINGp2 | Posterior Cingulate Gyrus (2) | M | -4 | -39 | 27 | 8 | -43 | 31 |
| f1_2 | superior frontal sulcus (2) | M | -27 | 56 | 1 | 28 | 56 | 7 |
| f2_2 | inferior frontal sulcus (2) | LM | -43 | 15 | 29 | 44 | 19 | 28 |
| F1_2 | Superior Frontal Gyrus (2) | L | -12 | 46 | 41 | 12 | 45 | 42 |
| F2_1 | Middle Frontal Gyrus (1) | M | -40 | 41 | 20 | 41 | 44 | 13 |
| F2O2 | Middle Frontal Gyrus: *Pars Orbitalis* (2) | M | -41 | 49 | -5 | 40 | 50 | -4 |
| F3O1 | Inferior Frontal Gyrus: *Pars Orbitalis* (1) | L | -42 | 31 | -17 | 44 | 33 | -14 |
| F3O2 | Inferior Frontal Gyrus: *Pars Orbitalis* (2) | M | -21 | 23 | -21 | 21 | 22 | -20 |
| F3t | Inferior Frontal Gyrus: *Pars Triangularis* (1) | L | -49 | 26 | 5 | 50 | 29 | 5 |
| FUS1 | Fusiform Gyrus (1) | M | -32 | -9 | -34 | 32 | -8 | -35 |
| HIPP1 | Hippocampal Gyrus (1) | M | -30 | -7 | -19 | 30 | -5 | -18 |
| HIPP2 | Hippocampal Gyrus (2) | M | -25 | -32 | -3 | 25 | -31 | -2 |
| INSa2 | Anterior Insula (2) | LM | -34 | 17 | -13 | 35 | 18 | -13 |
| INSa3 | Anterior Insula (3) | LM | -34 | 24 | 1 | 37 | 24 | 0 |
| INSa4 | Anterior Insula (4) | M | -41 | 15 | 3 | 41 | 15 | 4 |
| ips2 | intraparietal sulcus (2) | M | -34 | -58 | 46 | 37 | -52 | 48 |
| ips3 | intraparietal sulcus (3) | M | -27 | -60 | 44 | 26 | -62 | 46 |
| P2 | Inferior Parietal Gyrus (1) | M | -45 | -53 | 50 | 43 | -53 | 48 |
| pHIPP2 | Parahippocampal Gyrus (2) | M | -28 | -27 | -19 | 29 | -25 | -19 |
| prec1 | precentral sulcus (1) | M | -50 | 6 | 26 | 50 | 10 | 24 |
| prec4 | precentral sulcus (4) | LM | -42 | 1 | 50 | 44 | 1 | 48 |
| SMA2 | Supplementary Motor Area (2) | L | -11 | 18 | 63 | 11 | 18 | 63 |
| SMA3 | Supplementary Motor Area (3) | LM | -7 | 8 | 66 | 6 | 10 | 66 |
| SMG7 | Supramarginal Gyrus (7) | L | -55 | -52 | 26 | 55 | -46 | 33 |
| STS1 | superior temporal sulcus (1) | L | -50 | 14 | -22 | 52 | 13 | -26 |
| STS2 | superior temporal sulcus (2) | L | -55 | -7 | -13 | 54 | -2 | -15 |
| STS3 | superior temporal sulcus (3) | LM | -55 | -33 | -2 | 53 | -32 | 0 |
| STS4 | superior temporal sulcus (4) | L | -57 | -48 | 13 | 55 | -46 | 15 |
| T1_4 | Superior Temporal Gyrus (4) | L | -59 | -23 | 4 | 60 | -20 | 2 |
| T2_3 | Middle Temporal Gyrus (3) | LM | -61 | -35 | -5 | 62 | -31 | -5 |
| T2_4 | Middle Temporal Gyrus (4) | L | -53 | -59 | 7 | 57 | -53 | 3 |
| T3_3 | Inferior Temporal Gyrus (3) | M | -56 | -53 | -14 | 57 | -46 | -14 |
| T3_4 | Inferior Temporal Gyrus (4) | M | -50 | -61 | -8 | 54 | -58 | -11 |

The Language-and-Memory Network corresponds to 37 homotopic regions of interest^44^ (Fig. 1), either specialized for language^9^, episodic memory^54^, or both. Each region is described by its gradient asymmetry value. To identify regions with changing asymmetry across the lifespan, and as described by Roe and colleagues^45,46^, we used a factor-smooth Generalized Additive Mixed Model with Hemisphere×Age (*i*.*e*., age-related change in asymmetry) as the effect of interest.

Gradient significant age-related changes in asymmetry were found in 25 of the 37 regions of the Language-and-Memory Network (68% of the Language-and-Memory Network regions, all *p*_*FDR*_<0.024, Fig. 2). On the lateral surface of the temporal lobe, significant regions were localized alongside the superior temporal sulcus (STS1, STS2, STS3), extending to the Superior Temporal Gyrus dorsally (T1_4) and joining the posterior part of the Inferior Temporal Gyrus (T3_4) and ventrally, the Fusiform Gyrus (FUS4). Advancing toward the parietal lobe, the Supramarginal Gyrus (SMG7), the Inferior Parietal Gyrus (P2), and the intraparietal sulcus (ips3) also showed significant Hemisphere×Age interactions. On the lateral surface of the left frontal lobe, the regions showing a significant Hemisphere×Age interaction covered the pars triangularis part of the Inferior Frontal Gyrus (F3t), as well as the pars orbitalis (F2O2), the junction of the Middle Frontal Gyrus (F2_1) with the precentral sulcus (prec1, and prec4). The superior frontal sulcus (f1_2), the medial part of the Superior Frontal Gyrus (F1_2), and the pre-superior motor areas (SMA2 and SMA3) were also part of these areas in the frontal lobe. Three regions were located within the anterior Insula (INSa2, INSa3, and INSa4), while three others were located along the Hippocampal (HIPP1 and HIPP2) and paraHippocampal Gyri (pHIPP2). The Posterior Cingulum (CINGp2) was selected in the posterior medial wall using this approach. The 12 non-significant regions (all pFDR>0.174) were localized in the posterior part of the temporal (STS4, T2_3, T2_4, and T3_3) and the parietal lobes (AG1, AG2, and ips2), the anterior cingulate (CINGa2), the amygdala (AMYG), and the inferior frontal gyrus (F3_O1, F3_O2) and sulcus (f2_2).

**Figure 2.**
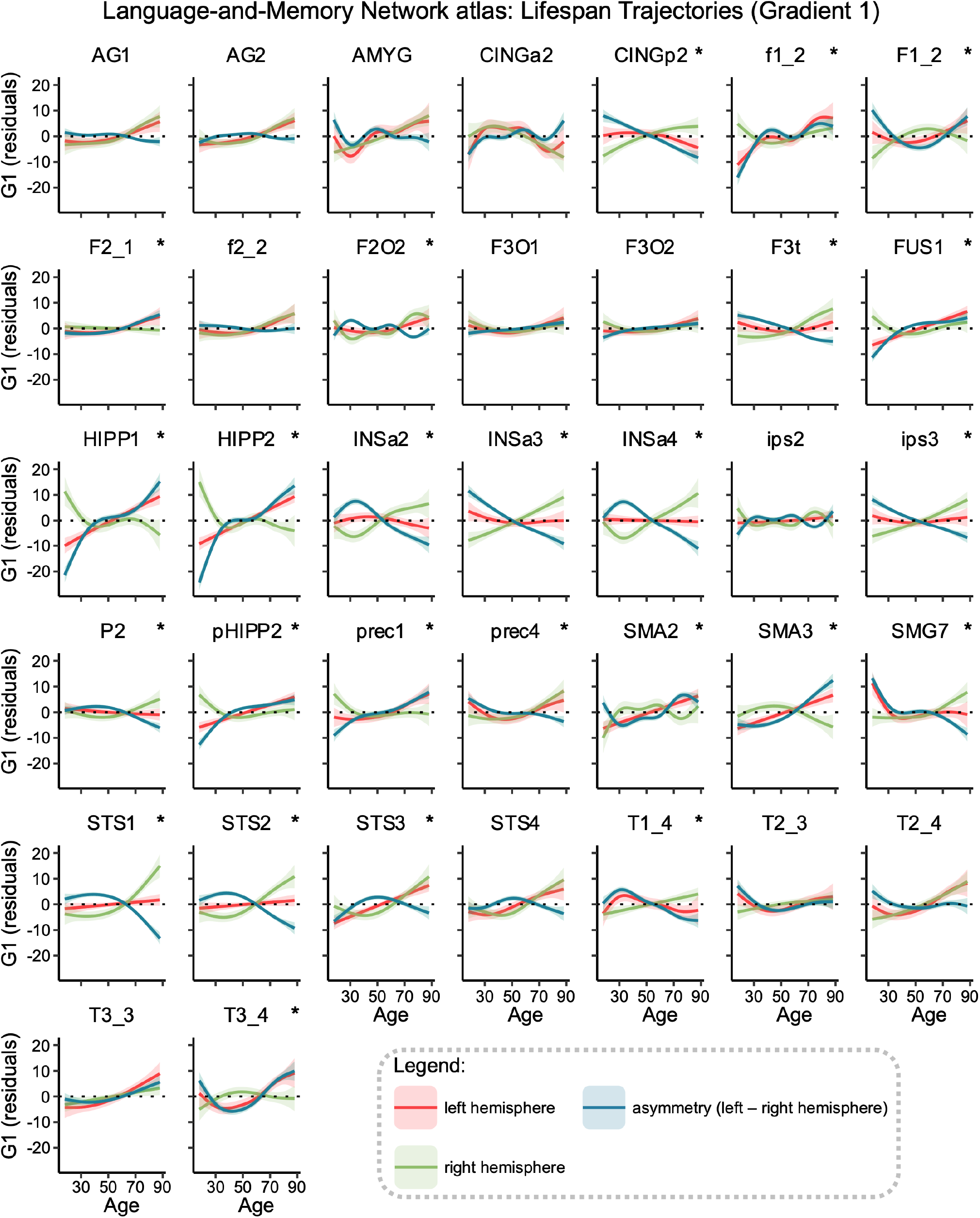
Gradient lifespan trajectories of Language-and-Memory regions. Each region’s graph shows the lifespan trajectory of the left (in red) and the right (in green) hemispheres and their asymmetry (in blue). Regions are plotted in alphabetical order. Trajectories were fitted using the generalized additive mixed models. Significant regions (*p*_*FDR*_<0.05) are marked with a star (*) in the top right corner. Data are residualized for sex, site, and random subject intercepts. Ribbons depict the standard error of the mean. The location of regions can be found in Fig. 1. Correspondences between the abbreviations and the full names of a region can be found in Table 1.

### Clustering of Asymmetry Trajectories

To investigate the asymmetry trajectories associated with the Hemisphere×Age interaction, we conducted clustering on the 25 significant regions within the Language-and-Memory Network to pinpoint areas displaying similar patterns of gradient asymmetry changes throughout adulthood (Fig. 2). The Partition Around Medoids algorithm identified two optimal partitions based on the mean silhouette width of 0.73. Including the regions that did not exhibit significant changes in gradient asymmetries over the lifespan, the Language-and-Memory Network regions are grouped into three distinct clusters (Fig. 3A).

**Figure 3.**
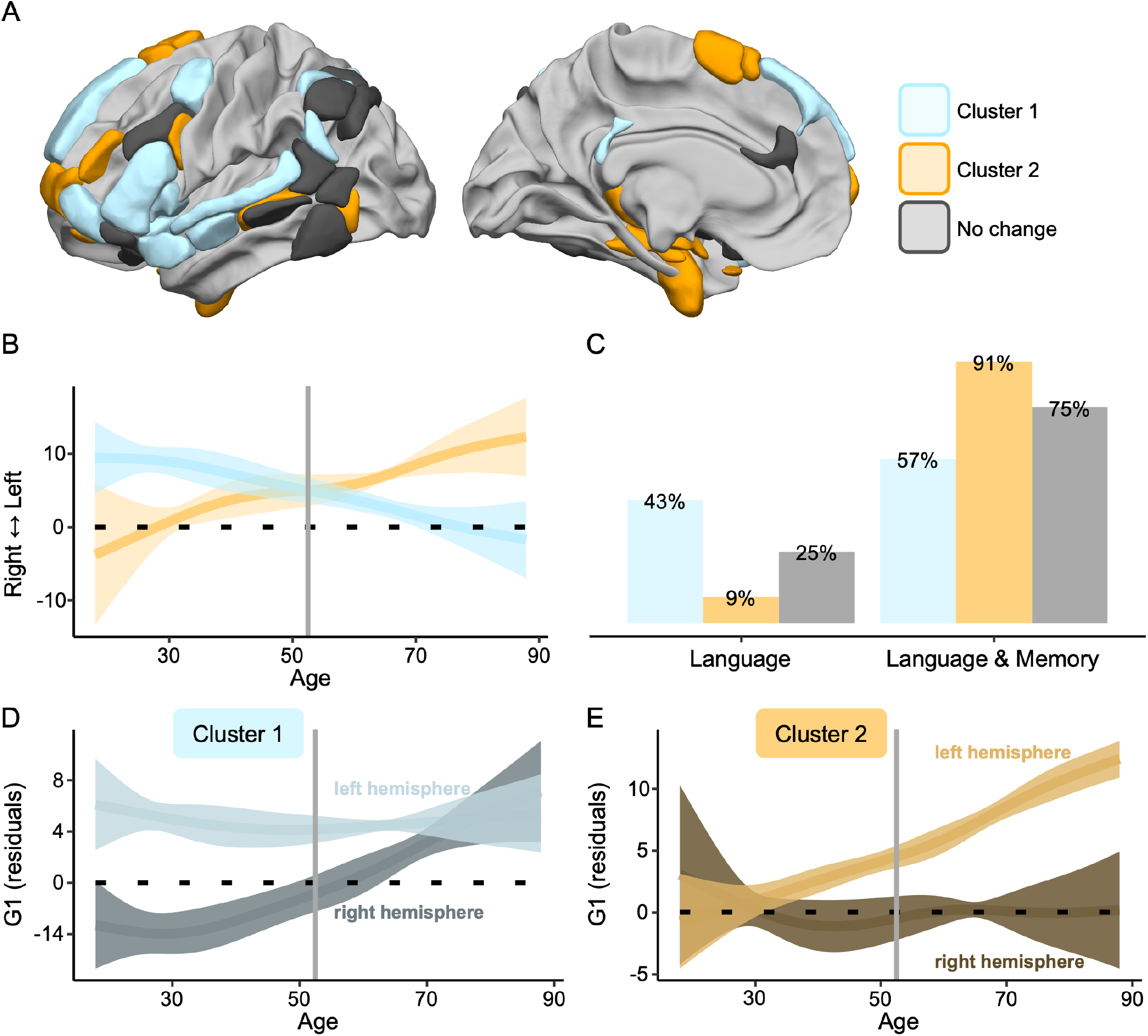
Patterns of language-related neurocognitive trajectories. **(A)** The 25 Language-and-Memory Network regions associated with the two main clusters of change, categorized according to the *k*-medoids classification applied to the Euclidean distance matrix derived from the age-related curves of asymmetry as modeled by the Generalized Additive Mixed Model. Cluster 1, in blue, changes from left-sided dominant to bilateral. Cluster 2, in orange, changes from a bilateral organization to a leftside dominance. See Fig. 1 and Table 1 for a description of the regions. **(B)** Average trajectory curves of the 1^st^ gradient asymmetries from 18 to 88 years old. The two main patterns of inverse changes (Cluster 1 and Cluster 2) with age. The vertical line represents the intersection point between Cluster 1 and Cluster 2: 52.55 years old, *i*.*e*., the age at which the 1^st^ gradient asymmetry trends reverse. Ribbons depict the standard deviation. **(C)** The proportion of each cluster depends on the underlying cognitive processes: language or language and memory. **(D-E)** Modeling of the average estimated 1^st^ gradient parameter for each hemisphere (left and right) across ages for Language-and-Memory Network regions belonging to Cluster 1 **(D)** and Cluster 2 **(E)**. Ribbons depict the standard deviation. The bilateralization of Cluster 1 with age is due to an increase of the 1^st^ gradient values in the right hemisphere, while the left hemisphere remains stable. The left-sided specialization of Cluster 2 with age is due to an increase of the 1^st^ gradient values in the left hemisphere, while the right hemisphere remains stable. This dual mechanism is mediated by an overspecialization of the contralateral hemisphere with age, characterized by an increased capacity to integrate high-level Language-and-Memory Network information.

The first cluster, highlighted in light blue in Fig. 3 and referenced similarly throughout the paper, comprised regions that showed an average increase in their gradient values in the right hemisphere (Fig. 3D). These regions transitioned to a slightly rightward asymmetrical state with aging (*smooth*_*88 yo*_=-1.72), whereas they exhibited leftward asymmetry in earlier life stages (*smooth*_*18 yo*_=9.40, negative slope from positive intercept, Fig. 3B). The right hemisphere heteromodality increased significantly with aging, while the left hemisphere capacity remained stable. Within this cluster, 43% of the regions were dedicated to processing language, while 57% were multimodal, handling language and memory functions (Fig. 3C). Cluster 1 regions are mapped onto the frontal, parietal, temporal, limbic cortices, and insula.

The second cluster, highlighted in light orange in Fig. 3 and referenced similarly throughout the paper, comprised regions that showed an average increase in their gradient values in the left hemisphere (Fig. 3E). These regions transitioned to a leftward asymmetry state with aging (*smooth*_*88 yo*_=12.23), whereas they exhibited rightward asymmetry organization in earlier life stages (*smooth*_*18 yo*_=-3.77, positive slope from negative intercept, Fig. 3B). The left hemisphere heteromodal specialization increased significantly with aging, while the right hemisphere capacity remained stable. Within this cluster, 9% of the regions were dedicated to processing language, while 91% were multimodal, handling language and memory functions (Fig. 3C). Cluster 2 regions are mapped onto the frontal, temporal, and limbic cortices.

The last cluster (in grey in Fig. 3), named “No change,” regrouped the 12 non-significant regions that showed no significant changes in their hemispheric asymmetries throughout the lifespan. This cluster encompasses 25% of regions exclusively associated with language function and 75% of the regions involved in language and memory processes.

The trajectories of clusters 1 and 2 indicated that the asymmetry switch occurred at 52.6 years old (Fig. 3B). From this age onward, Cluster 2, which mainly encompasses multimodal regions, became the dominant leftward asymmetrical cluster. Its heteromodality in later life surpassed the early life heteromodality of Cluster 1. Meanwhile, Cluster 1 continued its decline towards a symmetrical organization of information integration.

### Multimodal Brain-Cognition Association Change Analysis

Finally, we examined the extent to which changes in functional asymmetries among the two clusters are related to changes in language skills. To gain a comprehensive understanding, our analysis also incorporated the normalized volume of each region within the identified clusters. This approach allowed us to identify a tripartite relationship connecting anatomy, macroscale functional brain organization, and cognitive performance throughout aging. To achieve this, we used permutation-based Canonical Correlation Analysis (CCA) inference^55^. CCA reveals modes of joint variation between two sets of variables, resulting in a set of mutually uncorrelated modes. Each mode captures a portion of the multivariate brain and behavior covariation. The CCA was conducted between a set of brain variables (including gradients and normalized volumes) and a set of cognitive variables evaluating language performance (including naming and tip of the tongue for language production and accuracy and reaction time in language comprehension). Language skill assessments are described in the Methods section (Cognitive Assessment of Participants). Prior to conducting CCA, we summarized the high-dimensional set of brain variables using Principal Component Analysis^55^ (PCA). The CCA has been performed on the 554 participants of the CamCAN database only due to a lack of behavioral data for other participants. **Cluster 1 –** We first conducted a PCA on the brain set variables (gradient and normalized volume asymmetries) from the first cluster (Fig. 3A). This analysis indicated that the 28 variables could be condensed into four principal components, accounting for 49.79% of the total variance in the brain set. The first component alone explained 26.75% of the total variance and opposed the volume asymmetries of the dorsal language pathway regions to those of the ventral pathway regions (Fig. 4A, left column). Positive loadings then indicated a leftward asymmetry of the dorsal pathway, while negative ones indicated a rightward asymmetry of the ventral pathway. The second component alone explained 12.15% of the total variance. It opposed the volume asymmetries of the dorsal language pathway regions to those of the ventral pathway regions and the asymmetries of the first gradient (Fig. 4A, left column). Positive loadings then indicated a rightward asymmetry of the volume of the dorsal pathway regions and a leftward asymmetry of the ventral pathway as well as the gradient values. At the same time, negative loadings indicated the opposite pattern.

**Figure 4.**
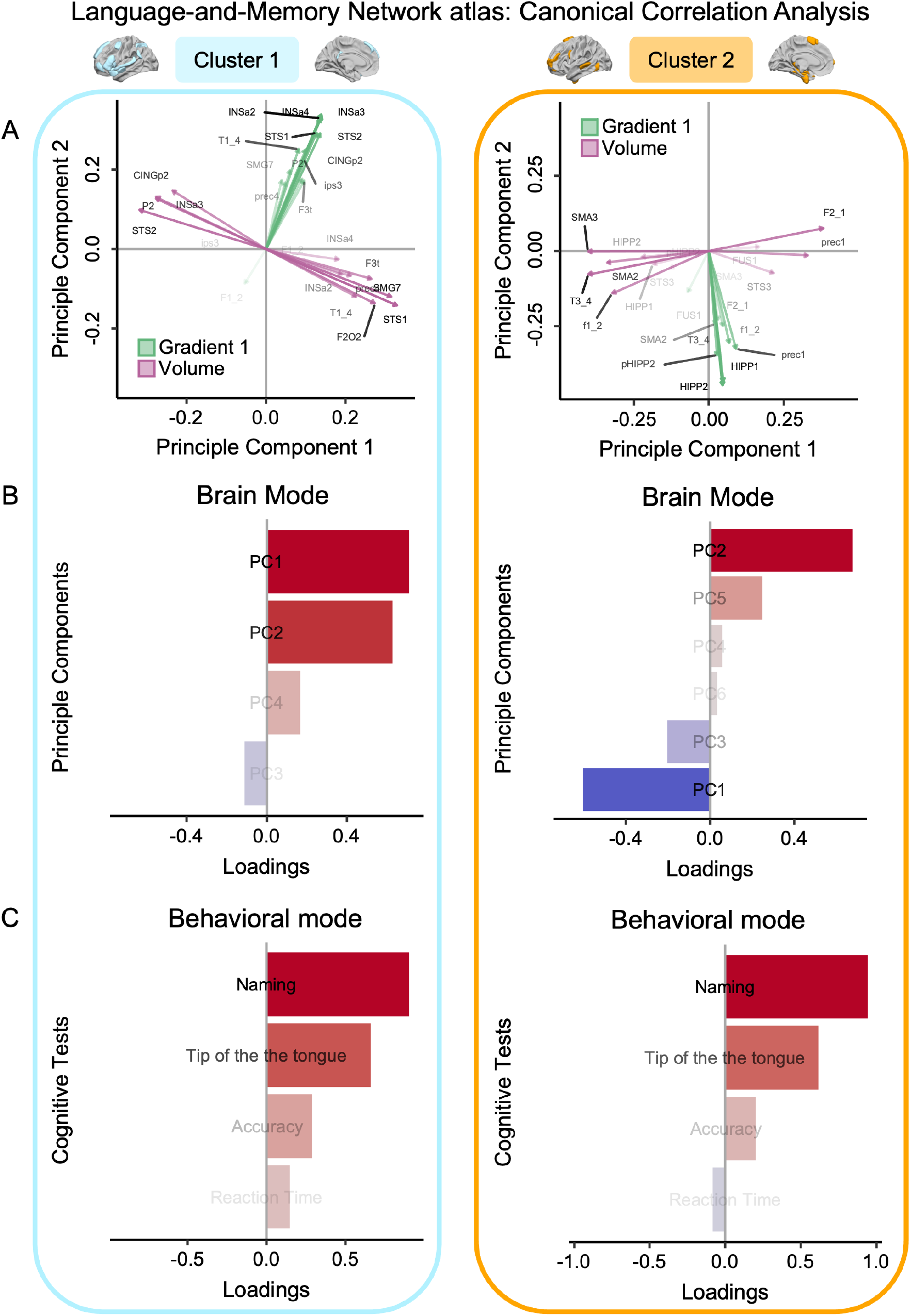
Brain-behavior association using canonical correlation analysis. **(A)** Biplot of the principal component analysis of the regions belonging to Cluster 1 (*n*=14, on the left) and Cluster 2 (*n*=11, on the right). Each region was characterized by its asymmetry values of the 1st gradient and normalized volume. The two principal components of Cluster 1 explained 38.89% of the total variance (*Principal Component 1*=26.74%, *Principal Component 2*=12.15%). The two principal components of Cluster 2 explained 33.34% of the total variance (*Principal Component 1*=20.47%, *Principal Component 2*=12.87%). For Cluster 1, the 1^st^ principal component opposed the volume asymmetries of the dorsal language pathway regions to the ventral semantic pathway regions. The 2^sd^ component opposed the symmetries of the 1st gradient to the symmetries of the normalized volume. For Cluster 2, the 1^st^ principal component opposed the asymmetry of mesial regions versus the volume asymmetry of lateral regions. The 2^sd^ component coded for the symmetry of the 1^st^ gradient, specifically, the symmetry of the temporo-mesial memory-related regions: a larger value meant a larger symmetry. **(B-C)** Overview of the canonical correlation analysis first modes. Only data from participants with all scores on the selected language indicators were included in the analysis (*n*=554; CamCAN cohort only). Sex, age, and general ⟵ cognitive status (MMSE) were entered as covariates. **(B)** First mode for brain variables. For Cluster 1, the brain mode explained 38% of the variance. It is saturated by the first two components of the principal component analysis, mixing the multimodal biomarkers included in the analysis (1^st^ gradient and normalized volume). For Cluster 2, the brain mode explained 24% of the variance. It is saturated by the first two components of the principal component analysis. **(C)** First mode of behavioral variables. For Cluster 1 and 2, the behavioral mode explained 39% of the variance and was saturated by the language production tasks involving lexical access and retrieval: naming and tip of the tongue. Results for Cluster 1 are framed in light blue. Results for Cluster 2 are framed in orange.

The multimodal canonical correlation analysis on the first cluster, which incorporated four brain metrics (principal components) and four behavioral metrics, revealed a single significant canonical correlation linking anatomy, function, and behavior (*p*_*FWER*_<1×10^−3^). This brain mode accounted for 37.58% of the variance and primarily reflected the first and second components of the brain data set (Fig. 4B, left column). Positive values of the brain mode were associated with positive loading values for both the first and second principal components. Specifically, these positive values in the brain mode indicated a leftward asymmetry for all regions regarding gradient and normalized volume in the dorsal language pathway regions. Conversely, they represented a rightward asymmetry in the ventral pathway regions. The behavioral mode accounted for 39.47% of the variance and primarily reflected the naming and tip of the tongue tests (Fig. 4C, left column). Positive values of the behavioral mode were associated with better performances in language production. The correlation between the brain and behavioral modes was 0.28, as depicted in Fig. 5 (left panel). Improved language production abilities were linked to a leftward asymmetry of the gradient value within the Language-and-Memory Network regions of the first cluster, a leftward asymmetry of the normalized volume for the dorsal language pathway regions, and a rightward asymmetry for the ventral language pathway regions.

**Figure 5.**
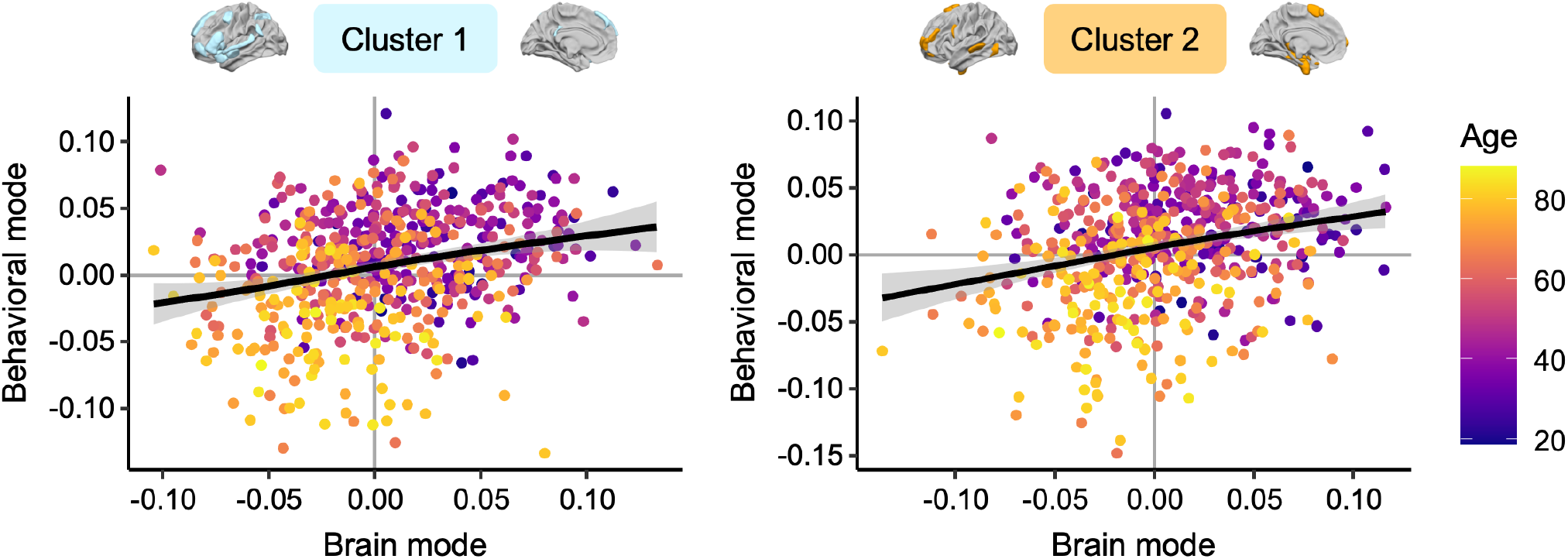
Relationship between changes in inter-hemispheric balances and their behavioral implications in a multimodal perspective. The first brain and behavioral modes were significantly correlated for both clusters: *r*=0.28, *p*<1.10^−3^. The significance of correlations between modes was assessed using permutation testing (*n*=1000). Color code for age.

#### Cluster 2

The principal components analysis on the brain set variables (22 variables, gradient, and normalized volume asymmetries) for the second cluster (Fig. 3A) resulted in six principal components. Together, these principal components explained 59.35% of the total variance in the brain set. The first component alone explained 20.48% of the total variance and opposed the volume asymmetries of the mesial regions to those of the lateral side (Fig. 4A, right column). Positive loadings then indicated a rightward asymmetry of the normalized volume of the mesial regions and a leftward asymmetry of the lateral regions. Negative loadings indicated the opposite pattern. The second component alone explained 12.86% of the total variance and captured the asymmetry of the gradient, specifically, the asymmetry of the temporo-mesial memory-related regions (Fig. 4A, right column). Positive loadings indicated a rightward asymmetry of the gradient, while negative loadings indicated a leftward asymmetry.

The multimodal canonical correlation analysis on the second cluster, which incorporated six brain metrics (principal components) and four behavioral metrics, revealed a single significant canonical correlation linking anatomy, function, and behavior (*p*_*FWER*_<1×10^−3^). This brain mode accounted for 23.61% of the variance and opposed the second component of the brain data set to the first one (Fig. 4B, right column). Positive values of the brain mode were associated with positive loading values for the second component and negative values for the first component. A positive brain mode value meant a leftward asymmetry of the normalized volume of the mesial regions, a rightward asymmetry of the lateral regions, and a rightward asymmetry of the gradient. The behavioral mode accounted for 39.04% of the variance and, similarly to Cluster 1, primarily reflected the naming and tip of the tongue tests (Fig. 4C, right column). The correlation between the brain and behavioral modes was 0.28, as depicted in Fig. 5 (right panel). Improved language production abilities were linked to a rightward asymmetry of the gradient value within the temporo-mesial memory-related regions, a leftward asymmetry of the normalized volume of the mesial regions, and a rightward asymmetry of the normalized volume of the lateral regions.

## Discussion

Our study uncovers that functional asymmetry in the integration of high-level information plays a pivotal role in the neural mechanisms underlying language processing and capabilities. Longitudinal analysis revealed shifts in hemispheric dominance, underscoring the dynamic nature of functional lateralization. These changes in asymmetry are associated with the language production challenges commonly seen in typical aging, disputing the notion that increased engagement of the contralateral hemisphere in older adults serves a compensatory role. Instead, our findings align with the brain maintenance theory, highlighting the importance of preserving a youthful functional brain state for optimal cognitive performance as individuals age. This study paves the way for further exploration into the dynamic processes by which the brain and cognition adapt throughout the aging process.

We found that this dual mechanism of the Language-and-Memory Network neurofunctional imbalance in integrating complex, high-level information begins after age 50 and intensifies over time (Fig. 4B). These findings are consistent with previous functional studies showing significant transitions in middle age^56^. They also align with the onset of structural changes observed in healthy older adults regarding cortical thickness asymmetry, showing an accelerated loss of asymmetry after midlife^46,57,58^. The reduction in structural asymmetry is notably significant in higher-order cortex and heteromodal regions, which may account for the extensive reorganization observed in the functional organization of the Language-and-Memory Network regions. None of these changes in asymmetry contributed to maintaining language performance with age and were, instead, linked to poorer performance. For Cluster 1 and Cluster 2, the pattern observed in young adults was related to more efficient language production (Fig. 6), underlining the importance of specialization at all ages for effective interhemispheric cooperation. Consequently, the changes do not support the hypothesis of a compensatory phenomenon^59^, preserving language performance with age. On the contrary, it aligns with the dedifferentiation theory of aging^60–63^ and the brain maintenance theory^64,65^, suggesting that maintaining a (functional) youthful brain state is essential to cognitive preservation as individuals age. These findings further underscore Roe and colleagues’ insights in their recent investigation of age-related shifts in functional asymmetry during memory retrieval^66^.

**Figure 6.**
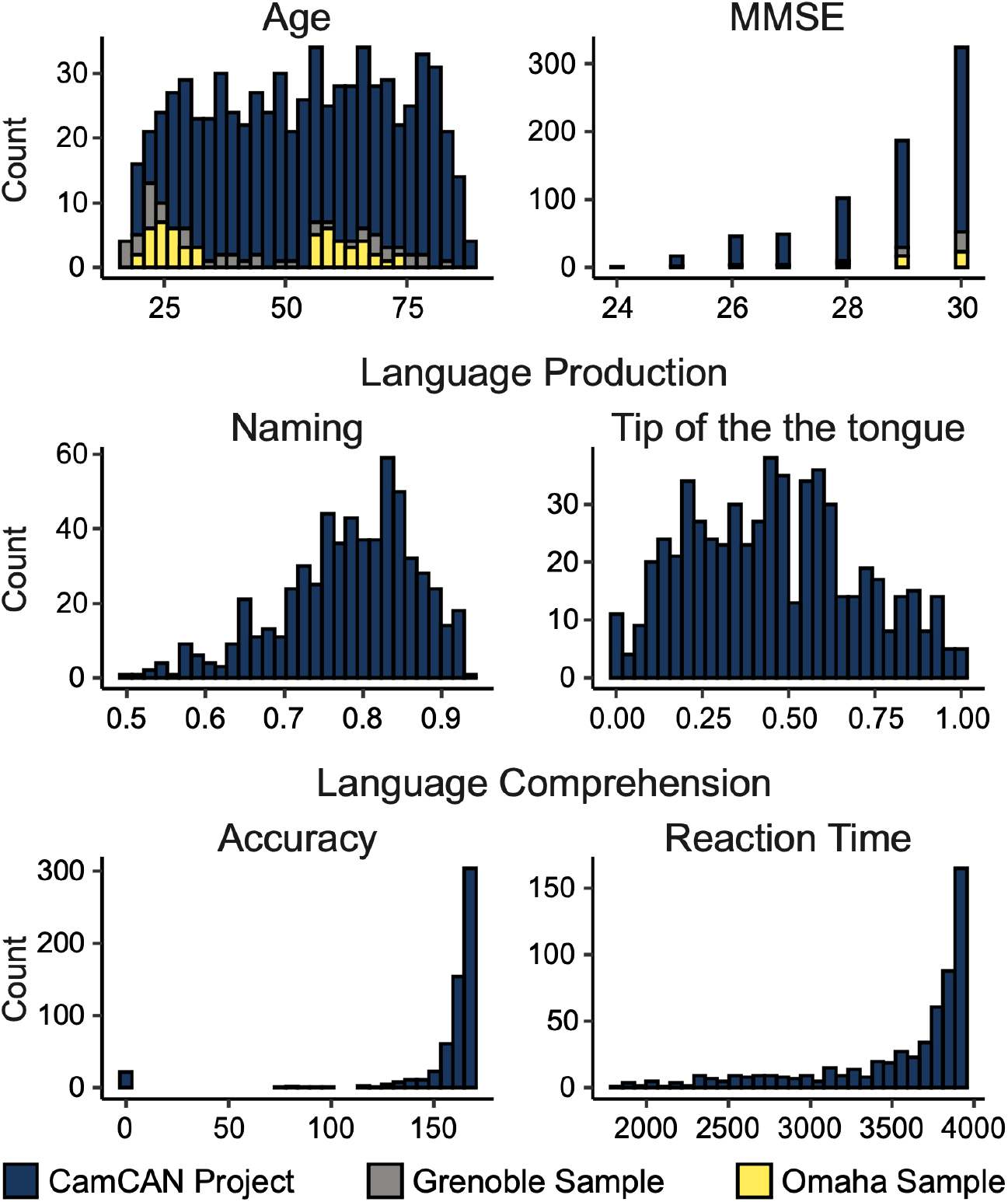
Age and behavioral performance stacked distributions. The behavioral tests assess various cognitive functions associated with language: word production, lexical access/retrieval abilities (picture Naming accuracy and Tip of the tongue ratio), and semantic and syntactic comprehension abilities (Accuracy and Reaction Time). A description of the behavioral variables is available as supplementary material in the article by West and colleagues^93^. Reaction Time and Tip of the tongue performance were inverted, so all scores close to zero represent worse performances. Stacked histograms for Age and MMSE include 728 participants. Language Production and Language Comprehension stacked histograms include 554 participants of the CamCAN database only due to a lack of behavioral data for other participants.

The lateralization of individual functions, such as language, may be closely associated with the lateralization of many seemingly independent processes^43^. Several studies suggest that the LH specialization for language may be linked to the concept of “complementary lateralization.” This stands in contrast to the preferential specialization of the contralateral hemisphere (the RH) for other high-level cognitive functions like visuospatial processing^67–71^. It has also been reported that the absence of functional lateralization for language production reduces performance in language tasks and other non-verbal, high-level functions^72^. The attentional and executive control networks^73^ play a role in maintaining these specializations, with LH control regions (Control-B) closer to the Default Mode Network (DMN-B) and RH attentional regions (DAN-B) nearer to the sensory-motor end of the gradient^38^. Importantly, control networks undergo extensive reconfigurations during the aging process^8,19,21,74–76^. These changes affect the substantial alterations observed in language’s functional asymmetries and other cognitive functions. Although beyond the scope of this research, studying how changes in neurofunctional equilibriums for different cognitive functions occur with age would offer invaluable insights into mutual network interactions.

The human brain typically exhibits marked structural left-right disparities, particularly pronounced in perisylvian regions associated with language. Although genetics contribute to these asymmetries, their impact appears to be less substantial than previously assumed, with heritability estimated at less than 30% in adults^4,77^, suggesting that environmental factors likely play a substantial role. Current research points to two primary developmental trajectories: the first is primarily influenced by genetics and lays the groundwork for brain lateralization, while the second, built upon this genetic foundation, entails prolonged development in brain regions responsible for complex functions, rendering them more susceptible to the influence of environmental factors^43^. The aging process, particularly affecting heteromodal associative brain regions in middle age, may introduce a phase of heightened vulnerability to environmental and life experience factors from this period onward. Pinpointing the specific environmental factors and midlife experiences that contribute to resilience or susceptibility in the face of changes in brain asymmetry holds the potential to enhance our understanding of the variability in neurocognitive aging. This may facilitate the development of personalized preventive measures and interventions for individuals at risk of experiencing accelerated aging. Importantly, functional asymmetry is not solely dependent on cognitive aspects but is also strongly influenced by sensory inputs^78,79^. The decline of the peripheral nervous system plays a pivotal role in triggering significant functional reconfigurations within the central nervous system^80,81^ for the consequences of age-related hearing loss on brain function. Furthermore, hearing impairment in midlife is a substantial risk factor for dementia, as emphasized in the 2020 report by the Lancet Commission on dementia prevention, intervention, and care^82^. By elucidating the intricate relationships between sensory inputs, neural adaptations, and cognitive aging, the investigation of bottom-up influences presents an intriguing yet relatively unexplored research avenue.

Several methodological considerations and potential biases require discussion. Our study isolated the effects of age from gender through statistical control. However, gender-based disparities in language-related functional connectivity have been reported^8^, alongside variations in the asymmetry of hemispheric functional gradients^40^. Hence, future studies must delve into gender-specific characteristics of the extended language network. Furthermore, given that certain aspects of brain aging manifest disparities between males and females^83^, special consideration should be given to older adults since gender differences could be amplified. Moreover, our study predominantly included participants from WEIRD (Western, Educated, Industrialized, Rich, and Democratic) societies. Considering that most of the global population does not fit within this category^84^, it would be beneficial to replicate these findings in more diverse populations, considering the importance of cultural diversity in research. Resting-state functional MRI has gained popularity due to its strong association with task-based fMRI activations^30,31^ and ease of acquisition, rendering it a valuable proxy for capturing functional neuronal processes. Nevertheless, the strength of hemispheric specialization for language depends on multiple factors, particularly the nature of the task^9,85^. Hence, conducting an additional study encompassing a diverse array of language-related functional tasks is essential to validate the consistency of the trends observed in our resting-state functional data. Open fMRI databases dedicated to language, such as InLang^8^, could facilitate such investigations. However, the databases available to date only sometimes include a wide age range, which could limit insights into older adults. Finally, longitudinal data are imperative for providing conclusive evidence regarding evolutionary trajectories throughout the lifespan and their cognitive implications. The STAC-r model (revised Scaffolding Theory of Aging and Cognition model) emphasizes the importance of examining cognitive changes within individuals^86^. This approach helps distinguish between mechanisms that maintain brain integrity and compensatory processes. Both mechanisms are crucial for preserving cognition in older adults, as noted by Reuter-Lorenz and Park in 2014. However, the current scarcity of extensive longitudinal cohorts, spanning both older and younger adults, hinders the identification of features predictive of future brain function and cognitive preservation^87^. It would also be important to extend the study to cohorts with mild cognitive impairment (MCI) and related conditions, which is crucial for assessing the specificity of the observed effects and discerning trends across different conditions.

## Methods

### Database Demographics

The study sample comprised three datasets, accumulating 728 healthy adults (371 women) from 18 to 88 years old (*μ*=52.84 years, SD=19.19 years, Fig. 6). Included participants had resting-state (rs) fMRI and structural MRI from a 3T scanner, meeting criteria of passing quality checks (*fmriprep* QC reports) and exhibiting no confirmed neurological or psychiatric pathologies.

The larger sample, the Cambridge Centre for Ageing and Neuroscience Project^88^ (CamCAN Project: www.mrccbu.cam.ac.uk), included 627 participants (316 women). Structural MRI data were acquired on a 3T Siemens TIM Trio scanner with a 32-channel head coil, using a T1-weighted, 3D MPRAGE sequence with the following parameters: Repetition Time (TR)/Echo Time (TE)/Inversion Time (TI)=2250/2.99/900*ms*, voxel size=1*mm* isotropic, flip angle=9°, Field of View (FOV)=256×240×192*mm*^*3*^, duration of acquisition: 4min 32s. For resting-state fMRI scans, participants rested with their eyes closed for 8*min* 40*s*. Two hundred and sixty-one brain volumes were acquired using a gradient echo planar imaging sequence (EPI, 32 axial slices, 3.7 *mm* thickness, TR=2.0*s*, TE=30*ms*, flip angle=78°, FOV=192×192*mm*^*2*^, voxel size=3×3×4.44*mm*^*3*^). Further recruitment information and the acquisition parameters have been described elsewhere^89^. The sample mean age was 54.28 years (SD=18.61 years). Participants’ handedness was defined based on the manual preference strength assessed with the Edinburgh inventory^90^: participants with a score below 30 were considered left-handers^91,92^, right-handers otherwise. The sample contained 56 left-handed participants (32 women). CamCAN funding was provided by the UK Biotechnology and Biological Sciences Research Council (grant number BB/H008217/1), with support from the UK Medical Research Council and the University of Cambridge, UK.

The second sample was collected in Omaha, NE, USA, and included 54 participants (31 women). The acquisition parameters are fully described in Doucet and colleagues^87^. Briefly, participants were scanned on a 3T Siemens Prisma scanner using a 64-channel head coil. Structural images were acquired using a T1-weighted, 3D magnetization-prepared rapid gradient-echo (MPRAGE) sequence with the following parameters: TR=2400*ms*, TE=2.22*ms*, FOV: 256×256*mm*, matrix size: 320×320, 0.8*mm* isotropic resolution, TI=1000*ms*, 8 degree-flip angle, band-width=220*Hz/Pixel*, echo spacing=7.5*ms*, in-plane acceleration GRAPPA (GeneRalized Autocalibrating Partial Parallel Acquisition) factor 2, total acquisition time ∼7*min*. Participants also completed a resting-state fMRI scan (eyes open). Scans were performed using a multi-band T2* sequence with the following acquisition parameters: TR=800*ms*, TE=37*ms*, voxel size=2×2×2*mm*^*3*^, echo spacing 0.58*ms*, bandwidth=2290*Hz/Pixel*, number of axial slices=72, multi-band acceleration factor=8, 460 volumes. The sample mean age was 44.13 years (SD=19.07 years). Participants’ handedness was self-reported: the sample contained seven left-handed participants (3 women). The Institutional Review Board for Research with Human Subjects approved the study at Boys Town National Research Hospital. Each participant provided written informed consent and completed the same protocol.

The third sample was collected in Grenoble, France, and included 47 participants (24 women). T1-weighted high-resolution three-dimensional anatomical volumes (T1TFE, 128 sagittal slices, 1.37*mm* thickness, FOV=224×256*mm*^*2*^, 0.8*mm* isotropic resolution) were acquired for each participant by using a whole-body 3 T MR Philips imager (Achieva 3.0 T TX Philips, Philips Medical Systems, Best, NL) with a 32-channel head coil. For resting-state fMRI scans, four hundred volumes were acquired using a gradient echo planar imaging sequence (FEEPI, 36 axial slices, 3.5*mm* thickness, TR=2.0*s*, TE=30*ms*, flip angle=75°, FOV=192×192*mm*^*2*^, voxel size=2×2×2*mm*^*3*^). Participants were asked to lie down in the scanner with eyes open on a central cross during the duration of the acquisition period (13*min* 20*s*). The sample mean age was 43.57 years (SD=21.92 years). Participants’ handedness was self-reported: the sample contained two left-handed participants (1 woman). The ethics committee of the Grenoble Alpes University Hospital approved data collection (CPP 09-CHUG-14; MS-14-102).

The *Supplementary Information* section (Comparative tables of database acquisition parameters) provides comparative tables of database acquisition parameters.

We used the whole age range of the sample (*n*=728, 18-88 years) to model the asymmetry trajectories further throughout the lifespan. By merging the CamCAN cohort with Grenoble and Omaha samples, we expanded our age coverage from 18 to 88 years old, addressing the lack of young adults in the CamCAN cohort (as depicted in Fig. 6), and making the age distribution uniform, allowing a more reliable analysis.

### Cognitive Assessment of Participants

For all 728 participants, we checked the Mini Mental State Examination (MMSE) scores to ensure that the general cognitive functioning of our sample remained within the expected range (*Q*_*1*_=28, *Q*_*3*_=30).

Among the three cohorts in our study, only the CamCAN cohort underwent an extensive set of behavioral assessments, resulting in cognitive data available for a specific sub-sample of 554 participants. These assessments, conducted outside the MRI scanner, are detailed in previous literature^89,94^. We limited our analyses to language skill assessments only (Fig. 6). We chose language-related measures because of their effectiveness in assessing diverse language-related aspects, encompassing word production, lexical access, and word retrieval (evaluated via picture naming accuracy and the tip-of-the-tongue ratio), as well as the understanding of semantics and syntax (measured through accuracy and reaction time). Further comprehensive descriptions of these behavioral variables are available in the supplementary materials provided by West and colleagues^93^.

### MRI Data Preprocessing

The neuroimaging data were formatted following the BIDS standard^44,95^ (Brain Imaging Data Structure - http://bids.neuroimaging.io/) and then preprocessed using the *fMRIPrep* software^96,97^ (https://fmriprep.org/en/stable/). The T1w preprocessing included skull stripping, tissue segmentation, and spatial normalization. Preprocessing of the rs-fMRI data followed the consensus steps for functional images, including motion correction, slice timing correction, susceptibility distortion correction, coregistration, and spatial normalization. The data were represented in the Montreal Neurological Institute (MNI) volumetric space. Finally, time series were extracted for each homotopic region of interest (described in the following subsection) using *Nilearn* (https://nilearn.github.io/) with nuisance parameter regression. Confounding regression included cerebrospinal fluid and white matter signals and translation and rotation parameters for *x, y*, and *z* directions.

### Language-and-Memory Network Statistics

Our statistical analyses were based on the Language-and-Memory Network atlas, an extended language network encompassing language-specific areas and related memory regions^44^. Briefly, the Language-and-Memory Network comprises 37 homotopic regions of interest. Among these 10 regions uniquely dedicated to the core supramodal language network^9^, 19 supporting episodic memory^54^, and eight regions underpinning both language and episodic memory processes. The core language network corresponded to a set of heteromodal brain regions significantly involved, leftward asymmetrical across three language contrasts (listening to, reading, and producing sentences), and functionally connected. The memory network was underpinned by areas that demonstrated strong activation patterns connected to episodic memory processes, such as encoding, effective recovery, and reminiscence. Fig. 1 shows the Language-and-Memory Network in a brain rendering, and Table 1 lists all the Language-and-Memory Network regions. It should be noted that the language atlas was based on the AICHA atlas, a functional brain homotopic atlas optimized for studying functional brain asymmetries^48^.

We computed two features characterizing the high-order Language-and-Memory Network regions^44^ from the preprocessed neuroimaging data: the normalized volume and the first functional gradient (G1) reflecting the macroscale functional organization of the cortex^37^. The first gradient captures the most variance of the correlations matrices (20%, 22%, and 19% for CamCAN, Omaha’s, and Grenoble’s cohorts, respectively). It has been previously shown to accurately reflect the lateralization of the language network^43^.

### Normalized Volume

Tissue segmentation was performed on the preprocessed T1w using the *FreeSurfer* pipeline (Version 6.0.0; CentOS Linux 6.10.i386). Briefly, the *FreeSurfer* segmentation process included the segmentation of the subcortical white matter and deep gray matter volumetric structures, intensity normalization, tessellation of the gray matter white matter boundary, automated topology correction, and surface deformation following intensity gradients to optimally place the gray/white and gray/cerebrospinal fluid borders at the location where the greatest shift in intensity defines the transition to the other tissue class. Structural volumes were normalized to total intracranial volume. Normalized volumes were extracted for each of the Language-and-Memory Network regions.

### Connectivity Embedding

Each participant’s values were obtained for the first functional gradient (G1). The gradients reflect participant connectivity matrices, reduced in their dimensionality through the approach of Margulies and colleagues^37^. Functional gradients reflect the topographical organization of the cortex in terms of sensory integration flow, as described by Mesulam^98^. Gradients were computed using Python (Python version 3.8.10) and the *BrainSpace* library^99^ (Python package version 0.1.3). Gradients computed at the regional and vertex levels performed similarly^99^.

Average region-level functional connectivity matrices were generated for each individual across the entire cortex (*i*.*e*., 384 AICHA brain regions). Consistent with prior work, each region’s top 10% connections were retained, and other elements in the matrix were set to 0 to enforce sparsity^37,49^. The normalized angle distance between any two rows of a matrix was calculated to obtain a symmetrical similarity matrix. Diffusion map embedding^50,100,101^ was implemented on the similarity matrix to derive the first gradient. Note that the individual-level gradients were aligned using Procrustes rotation (*N*_*iterations*_=10) to the corresponding group-level gradient. This alignment procedure was used to improve the similarity of the individual-level gradients to those from prior literature. Minmax normalization (0-100) was performed at the individual level for the whole brain^38^.

Gradient asymmetry was then computed for each participant and region. For a given region, gradient asymmetry corresponded to the difference between the normalized gradient value in the left hemisphere minus the gradient values in the right hemisphere. A positive gradient asymmetry value meant a leftward asymmetry; a negative value meant a rightward asymmetry.

### Statistical Analyses

Statistical analysis was performed using R^102^ (R version 4.2.2). Data wrangling was performed using the R library *dplyr*^103^ (R package version 1.0.10). Graphs were realized using the R library *ggplot2*^104^ (R package version 3.4.2). Brain visualizations were realized using Surf Ice^105^.

#### Modeling Gradient Asymmetry Trajectories Throughout Life

For each region of the Language-and-Memory Network, we used factor-smooth Generalized Additive Mixed Models (GAMMs, as implemented in the R library *gamm4*^106^; R package version 0.2-6) to fit a smooth gradient trajectory for Age per Hemisphere^45,46^ and to assess the smooth interaction between Hemisphere×Age within the clusters (see clusters definition below). Hemisphere was included as a fixed effect, while Sex and Site were treated as covariates of no interest. A random intercept for each subject was also included. GAMMs leverage smooth functions to model the non-linear trajectories of mean levels across individuals, providing robust estimates that can be applied to cross-sectional and longitudinal cognitive data^107^. GAMMs were implemented using splines, a series of polynomial functions joined together at specific points, known as knots. The splines allow the smooth function to adapt its shape flexibly to the underlying pattern in the data across the range of the predictor variable. This connection allows for the modeling of complex, non-linear relationships piecewise while maintaining continuity and smoothness across the function. To minimize overfitting, the number of knots was constrained to be low (*k*=6). The significance of the smooth Hemisphere×Age interaction was assessed by testing for a difference in the smooth term of Age between hemispheres. We applied a False Discovery Rate correction^108^ (FDR) to control for the number of tests conducted. Lastly, we used the linear predictor matrix of the GAMMs to obtain asymmetry trajectories underlying the interaction Hemisphere×Age and their confidence intervals. These were computed as the difference between zero-centered (*i*.*e*., demeaned) hemispheric age trajectories.

#### Classification of Age-Asymmetry Trajectories

To classify the regions of the Language-and-Memory Network found significant (after applying the FDR correction) according to their functional asymmetry skewness profile (*i*.*e*., increasing leftward asymmetry from baseline, decreasing leftward asymmetry, or stabilizing asymmetry with age), we computed a dissimilarity matrix (sum of square differences) between all trajectories. We applied the Partition Around Medoids algorithm (R library *cluster*^109^; R package version 2.1.4) to identify clusters of regions sharing identical lifespan trajectories. Clustering solutions from two to seven were considered, and the mean silhouette width determined the optimal solution.

#### Canonical Correlation Analysis to Assess Brain–Behavior Associations

For each cluster, we assessed the linear relationship between the gradient asymmetry trajectories of the Language-and-Memory Network, their normalized volume, and cognitive language performance using permutation-based Canonical Correlation Analyses^55^ (CCA) inference. CCA is a multivariate statistical method identifying linear combinations of two sets of variables that correlate maximally. CCA reveals modes of joint variation, shedding light on the relationship between cognitive language performance (behavioral set), the lifespan trajectories of sensory integration flow asymmetry, and its underlying anatomy (brain set). The CCA results on a set of *m* mutually uncorrelated (*i*.*e*., orthogonal) modes. Each mode captures a unique fraction of the multivariate brain and behavior covariation that isn’t explained by any of the other *m*−1 modes. To assess statistical significance, we determined the robustness of each estimated CCA mode using permutation testing with 1,000 permutations. This test computes *p*-values to assess the null hypothesis of no correlation between components, adhering to the resampling method developed by Winker and colleagues^110^. *p*-values were controlled over Family-Wise Error Rate (FWER; FWER corrected *p*-values are denoted *p*_*FWER*_), which is more appropriate than the FDR correction when measuring the significant canonical modes^110^.

Before conducting the CCA, we summarized the high-dimensional set of brain variables (gradient and normalized volume asymmetries) using principal component analysis^55^ (PCA). We retained components corresponding to the elbow point in the curve, representing the variance explained by each successive principal component. This was achieved using the R library *PCAtools*^111^ (R package version 2.5.15). These retained principal components were then designated as the brain set for the CCA. Finally, we residualized the two variable sets (brain and behavior sets) to remove the influence of sex, age, and MMSE before executing the CCA.

The CCA has only been realized on the 554 participants of the CamCAN database due to a lack of behavioral data for other participants.

## Data and Code Availability Statement

The CamCAN database is available upon request^88,89^: https://www.cam-can.org/. The second database collected in Omaha (NE, USA) is available on request from G.E.D. The third database collected in Grenoble (France) is available on request from M.B. These databases (collected in Omaha and Grenoble) are not publicly available due to privacy or ethical restrictions.

The atlas and the code used to produce the results and visualizations can be found here^112^: https://github.com/loiclabache/RogerLabache_2023_LanguAging.

## CRediT Authorship Contribution Statement

**Elise Roger:** Conceptualization, Data curation, Formal analysis, Investigation, Methodology, Software, Validation, Visualization, Writing - original draft, Writing - review & editing. **Loïc Labache:** Conceptualization, Data curation, Formal analysis, Methodology, Software, Validation, Visualization, Writing - original draft, Writing - review & editing. **Noah Hamlin:** Data curation, Investigation. **Jordanna Kruse:** Data curation, Investigation. **Monica Baciu:** Conceptualization, Data curation, Funding acquisition, Investigation, Methodology, Resources, Supervision, Writing - review & editing. **Gaelle E. Doucet:** Conceptualization, Data curation, Funding acquisition, Investigation, Supervision, Writing - review & editing.

## Supplementary Information

### Comparative tables of database acquisition parameters

#### MRI scanner

CamCAN: 3T Siemens TIM Trio scanner with a 32-channel head coil

Omaha: 3T Siemens Prisma scanner with a 64-channel head coil

Grenoble: 3T Philips Achieva TX scanner with a 32-channel head coil

#### Scan type: Structural MRI scans (T1-weighted)

| Cohort | Sequence | TR<br>(ms) | TE<br>(ms) | Flip angle<br>(°) | FOV<br>(mm) | Voxel size<br>(mm) | Other |
| --- | --- | --- | --- | --- | --- | --- | --- |
| CamCAN | MPRAGE | 2250 | 2.99 | 9 | 256 × 240 × 192 | 1 × 1 × 1 | GRAPPA: 2; TI: 900 ms |
| Omaha | MPRAGE | 2400 | 2.22 | 8 | 256 × 256 × 256 | 0.8 × 0.8 × 0.8 | GRAPPA: 2; TI: 1000 ms |
| Grenoble | MPRAGE | 2100 | 2.36 | 8 | 256 × 240 × 160 | 0.8 × 0.8 × 0.8 | GRAPPA: 2; TI: 1000 ms |

Notes. TR = repetition time; TE = echo time; FOV = field of view; MPRAGE = magnetization, prepared gradient echo; GRAPPA = GeneRalized Autocalibrating Partial Parallel Acquisition; TI = inversion time.

#### Scan type: Functional MRI scans (resting-state)

| Cohort | Sequence | TR<br>(ms) | TE<br>(ms) | Flip angle (°) | FOV<br>(mm) | Voxel size<br>(mm) | Volumes<br>(N) | Slices<br>(N) | Duration | Task |
| --- | --- | --- | --- | --- | --- | --- | --- | --- | --- | --- |
| CamCAN | EPI | 1970 | 30 | 78 | 192×192 | 3×3×4.44 | 261 | 32 | 8min 40s | Rest with eyes closed |
| Omaha | Multi-band EPI | 800 | 37 | 78 | 192×192 | 2×2×2 | 460 | 72 | 11min 83s | Rest with eyes on a fixation cross |
| Grenoble | EPI | 2000 | 30 | 75 | 192×192 | 2×2×2 | 400 | 36 | 13min 20s | Rest with eyes on a fixation cross |

Notes. TR = repetition time; TE = echo time; FOV = field of view; EPI = T2\*-weighted gradient echo planar image.

